# Dynamin-2 is a fission factor for SNX1-coated endosomal tubules

**DOI:** 10.64898/2026.10.16.759699

**Authors:** Meghadeepa Sarkar, Sannidhya De, Thomas J. Pucadyil

## Abstract

Sorting nexins of the SNX-BAR family shape endosomal membranes into tubular carriers that route cargo away from the degradative pathway, but we poorly understand the machinery that cuts these tubules to release transport carriers. Here we show that Dynamin-2 (Dyn2) is a fission factor for SNX1-coated tubules. In HeLa cells, Dyn2 localises to SNX1 tubules, and its knockout lengthens them and produces interconnected networks. Dyn2 binds SNX1 directly, and this interaction requires both the proline-rich domain of Dyn2 and the N-terminus of SNX1. On supported lipid bilayers, SNX1 alone tubulates membranes, and adding Dyn2 and GTP converts the emergent tubules into vesicles, indicating that no additional factors are needed for this activity. In cells, loss of Dyn2 redirects the retromer cargo GLUT1 to a LAMP1-positive compartment and reduces glucose uptake. These findings identify Dyn2 as the enzyme that releases SNX1 carriers and link this scission step to cargo recycling.

## Introduction

Endosomes sort membrane proteins between lysosomal degradation and return to the cell surface or the Golgi. Retromer and its associated sorting nexins (SNXs) capture cargo into tubular carriers that bud from the endosome and are released for transport (Burd and Cullen, 2014; Cullen and Steinberg, 2018). In SNX27–retromer, the cargo-selective VPS26–VPS29–VPS35 trimer works with SNX27, which recognizes PDZ-binding motifs such as the one in GLUT1 (Wieman et al., 2009; Steinberg et al., 2013). In ESCPE-1, SNX1/2–SNX5/6 heterodimers mediate retrograde transport of cargo such as CI-MPR independently of the trimer (Kvainickas et al., 2017; Simonetti et al., 2017). When these pathways fail, cargo is diverted to lysosomes. Retromer dysfunction is linked to Parkinson disease, Alzheimer disease and other neurological disorders (Small et al., 2005; Vilariño-Güell et al., 2011; Zimprich et al., 2011; Small and Petsko, 2015), and is exploited by pathogens to avoid reaching the lysosome (Finsel et al., 2013; Popa et al., 2015; Lu et al., 2022).

SNX1 and SNX2 are BAR-domain proteins that bind phosphoinositide-containing membranes and form the tubular coat of these carriers. Their disordered N-termini act as assembly hubs, with Asp-Leu-Phe (DLF) motifs present in these regions engaging the SNX27 FERM domain and the same regions binding VPS29 weakly (Yong et al., 2021; Shortill, Frier and Conibear, 2022; Simonetti et al., 2022). Although we understand cargo sorting and tubule formation reasonably well, how cells cut tubules to release carriers remains unclear. We think scission requires the coordinated action of several components, but the details remain to be defined (Cullen and Steinberg, 2018). Proposed mechanisms include actin-based and motor-based forces. The WASH complex drives Arp2/3-dependent actin polymerization on endosomes, and its loss produces long tubules that motors pull along microtubules (Derivery et al., 2009; Gomez and Billadeau, 2009). Pulling by dynein-dynactin may generate tension that favors release (Wassmer et al., 2009). We have also long suspected a role for dynamin. Dominant-negative dynamin mutants caused CI-MPR-containing tubules to accumulate (Nicoziani et al., 2000), and WASH co-precipitates with dynamin-2 (Dyn2) (Derivery et al., 2009). However, these studies relied on dominant-negative mutants or the inhibitor dynasore, and none linked dynamin to a specific tubule-forming coat. Work in yeast has made this link more concrete. The dynamin-related protein Vps1 localizes to endosomes decorated with retromer SNX-BARs, and its loss expands the coated tubular network and sends the cargo receptor Vps10 to the vacuole (Chi et al., 2014). The SNX-BAR Mvp1 binds Vps1 and recruits it to sever recycling tubules (Suzuki et al., 2021), and Vps1 alone severs phosphoinositide-containing tubules in a GTP-dependent manner in vitro (Gopan et al., 2026). Whether mammalian dynamin acts directly on SNX1-coated tubules has not been tested.

Here we address this question and show that Dyn2 is a fission factor for SNX1-coated endosomal tubules. In HeLa cells, Dyn2 localizes to SNX1 tubules, and its knockout lengthens them. Dyn2 binds SNX1 directly, an interaction that depends largely on its proline-rich domain and requires the SNX1 N-terminus. On supported lipid bilayers, SNX1, Dyn2 and GTP are sufficient to convert tubules into vesicles. Loss of Dyn2 also redirects GLUT1 to a LAMP1-positive compartment and lowers glucose uptake. Together, these results identify Dyn2 as a fission enzyme for SNX1 carriers and link this step to cargo recycling.

## Results

### Dyn2 localizes to SNX1-positive tubules and restricts their length in cells

To test whether Dyn2 is positioned to act on SNX1-coated membranes, we imaged HeLa cells expressing GFP-SNX1 and immunostained for endogenous Dyn2. GFP-SNX1 decorated both punctate endosomal structures and extended tubules. Dyn2 appeared as discrete puncta, and a subset of these puncta lay along GFP-SNX1 tubules (Fig. 1A, arrowheads in insets). We then asked whether Dyn2 influences SNX1 tubule morphology. We used a Dyn2 knockout (KO) HeLa line, in which immunoblotting confirmed complete loss of Dyn2 (Fig. 1B). To rule out expression differences as a source of morphological variation, we stably expressed GFP-SNX1 in WT and Dyn2 KO cells and FACS-sorted both lines for low and comparable GFP-SNX1 levels. Immunoblotting with an anti-GFP antibody confirmed that the levels matched (Fig. 1C). In WT cells, GFP-SNX1 was mostly punctate or vesicular, with only occasional short tubules. Dyn2 KO cells instead showed more abundant and visibly longer SNX1 tubules, which often formed extended, interconnected networks (Fig. 1D). Quantification of the longest SNX1 tubule per cell confirmed this (Fig. 1E). The mean of the experiment medians was 7.5 µm (SD = 0.6) in WT cells and 15.7 µm (SD = 5.9) in Dyn2 KO cells, roughly a two-fold increase (P < 0.0001, two-sided Mann-Whitney U test; n = 195 WT and 85 KO cells from three independent experiments). Dyn2 KO cells therefore accumulate elongated SNX1 tubules, which suggests that Dyn2 normally limits tubule length, possibly by severing them.

**Fig. 1.**
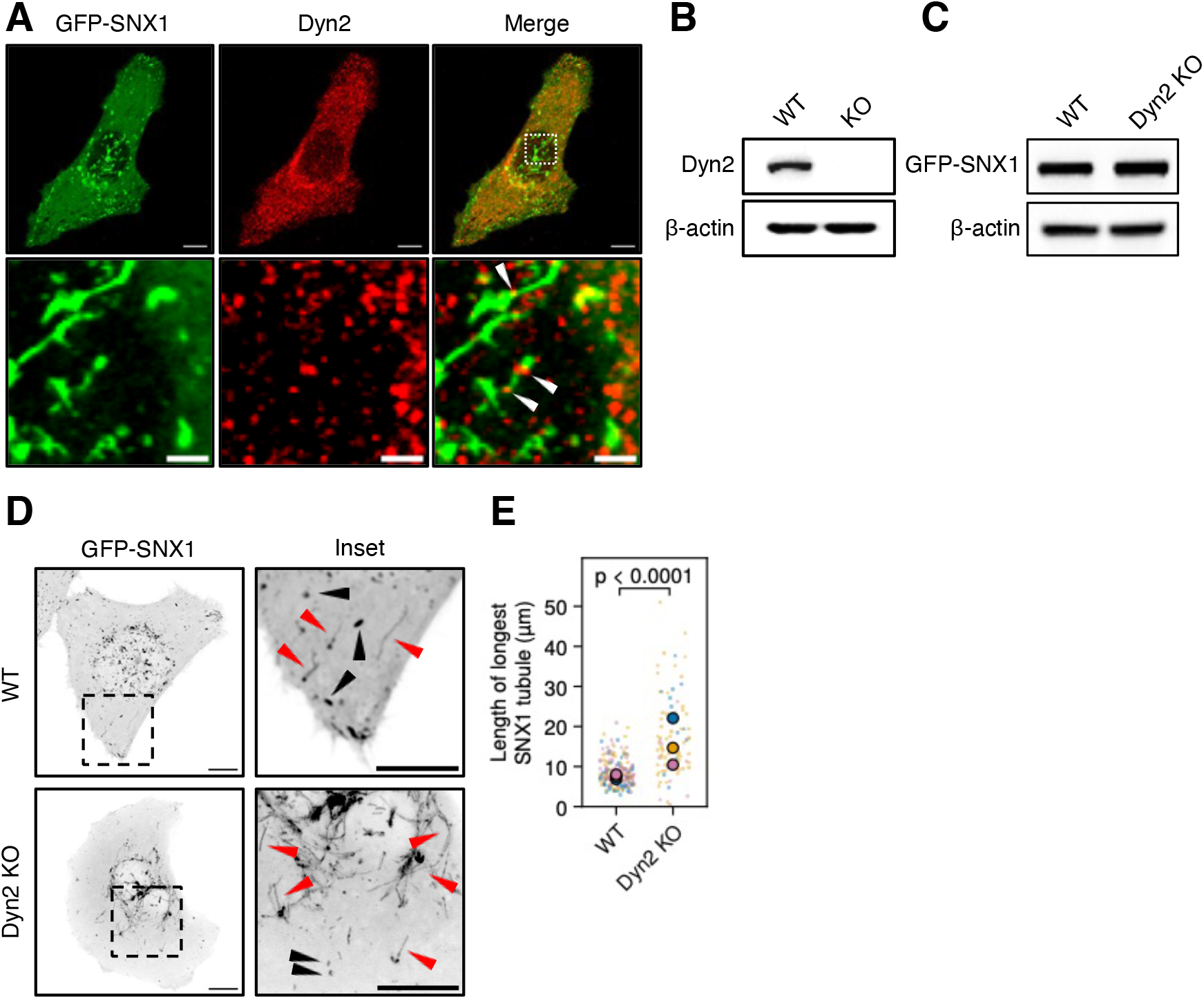
Dyn2 localizes to SNX1-positive tubules, and its loss leads to elongated SNX1 tubules. (A) Immunofluorescence of cells expressing GFP-SNX1 (green) and stained for endogenous Dyn2 (red). The boxed region in the merged image is magnified in the insets below. White arrowheads mark Dyn2 puncta on GFP-SNX1 tubules. Scale bars, 10 µm (top row) and 2 µm (insets). n > 30 cells imaged from one experiment. (B) Immunoblot for Dyn2 in lysates from HeLa WT and Dyn2 knockout (KO) cells, confirming loss of Dyn2 in the KO line. β-actin is the loading control. (C) Immunoblot for GFP-SNX1 in lysates from WT and Dyn2 KO HeLa cells stably expressing GFP-SNX1, showing comparable expression in both lines. β-actin is the loading control. Blots in B and C are representative of three independent experiments. (D) Confocal maximum intensity projections (MIPs) of WT and Dyn2 KO cells stably expressing GFP-SNX1 (inverted grayscale). Dashed boxes mark the regions magnified in the insets (right). Red arrowheads indicate SNX1 tubules, and black arrowheads indicate punctate or vesicular SNX1 compartments. Dyn2 KO cells show more abundant and longer SNX1 tubules than WT cells. Scale bars, 10 µm (whole cell) and 10 µm (insets). (E) Length of the longest SNX1 tubule per cell in WT and Dyn2 KO cells. Small dots represent individual cells, and large dots represent the median of each independent experiment (color-coded by experiment). n = 195 WT and 85 Dyn2 KO cells from three independent experiments. A two-sided Mann-Whitney U test was used for statistical comparison.

### Proline-rich domain of Dyn2 directly interacts with the N-terminus of SNX1

We next asked whether Dyn2 and SNX1 associate physically. Anti-GFP immunoprecipitation efficiently recovered both GFP-SNX1 and GFP alone (Fig. 2A). Endogenous Dyn2 co-precipitated with GFP-SNX1 but not with GFP (Fig. 2B), indicating a specific association in cells. To test whether the interaction is direct and to map its determinants, we used a bead-based binding assay with purified proteins. Dyn2-mCherry (CH) was strongly recruited to beads coated with GFP-SNX1 but not to beads coated with GFP, and GFP-SNX1 did not appreciably bind CH-coated control beads (Fig. 2C, D). Deleting the proline-rich domain (Dyn2ΔPRD-CH) reduced recruitment to GFP-SNX1 beads about 15-fold (median 0.08 versus 1.2, normalized to the signal on CH-bait beads; p < 0.0001, Mann-Whitney U test) (Fig. 2C, D). Conversely, beads coated with a SNX1 construct lacking its N-terminal region (GFP-SNX1ΔN) failed to recruit appreciable levels of either Dyn2-CH or Dyn2ΔPRD-CH (Fig. 2C, D). Together, these data show that Dyn2 binds SNX1 directly and that this binding depends largely on the Dyn2 PRD and requires the N-terminal region of SNX1.

**Fig. 2.**
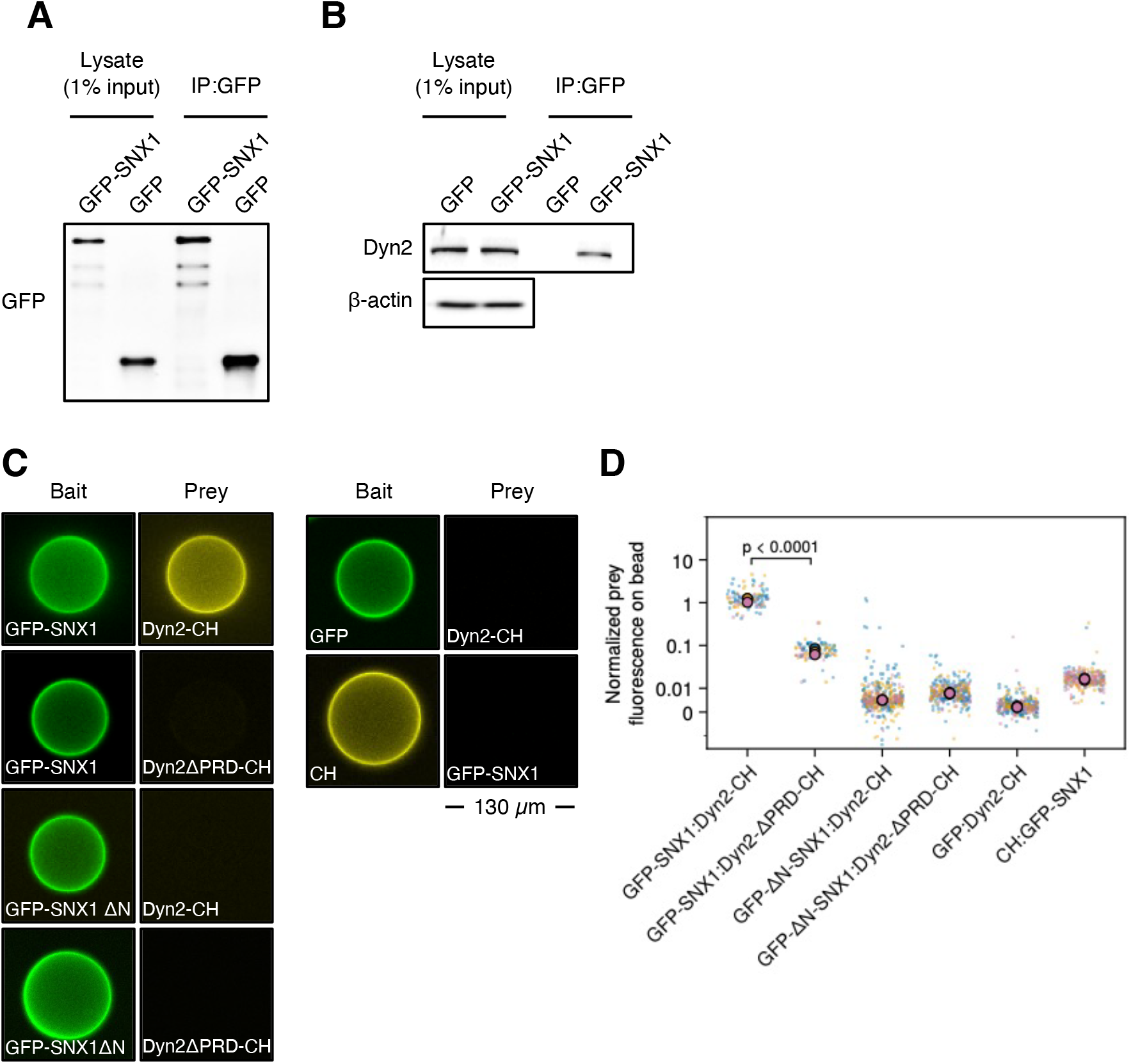
The proline-rich domain of Dyn2 directly interacts with the N-terminus of SNX1. (A) Anti-GFP immunoblot of lysates (1% input; left) and anti-GFP immunoprecipitates (IP:GFP; right) from cells expressing GFP-SNX1 or GFP, confirming expression and recovery of both proteins. (B) Immunoblot for endogenous Dyn2 in lysates (1% input; left) and anti-GFP immunoprecipitates (right) from cells expressing GFP or GFP-SNX1. β-actin is the loading control for the lysates. Dyn2 co-precipitated with GFP-SNX1 but not with GFP. Blots in A and B are representative of three independent experiments. (C) Bead-based binding assay with purified proteins. In each pair of panels, the bait coating the beads is shown on the left and the prey on the right. Scale bar, 130 µm. (D) Quantification of bead-based binding. X-axis represents the bait:prey combinations. The signal is normalized to GFP-bait or CH-bait no-prey beads. Small dots are individual beads and large circles are medians of independent sets, coloured by set. A two-sided Mann-Whitney U test was used for statistical comparison. Pooled bead numbers (n) and numbers of independent sets (N), GFP-SNX1:Dyn2-CH, n = 81, N = 3; GFP-SNX1:Dyn2-ΔPRD-CH, n = 67, N = 3; GFP-ΔN-SNX1:Dyn2-CH, n = 221, N = 3; GFP-ΔN-SNX1:Dyn2-ΔPRD-CH, n = 172, N = 3; GFP:Dyn2-CH, n = 168, N = 3; CH:GFP-SNX1, n = 152, N = 3.

### Dyn2 severs SNX1-generated membrane tubules in vitro

The elongated tubules in Dyn2 KO cells suggest that Dyn2 acts as a fission factor for SNX1 tubules. To test this in a defined system, we reconstituted tubulation on supported lipid bilayers (SLBs). When we flowed GFP-SNX1 alone onto an SLB, it progressively tubulated the membrane and produced an extensive network of tubules (Fig. 3A). The tubules contained the lipid probe, confirming that they are membrane tubules (Fig. 3A, right). Fluorescence recovery after photobleaching (FRAP) of the lipid probe showed recovery of the bleached membrane signal (Fig.3B), indicating that these tubules remain connected to the underlying bilayer. When we flowed GFP-SNX1 onto SLBs together with Dyn2 and GTP, tubules did not accumulate. Instead, GFP-SNX1 concentrated in discrete puncta whose number increased over time (Fig. 3C). These puncta contained the lipid probe, GFP-SNX1 and Dyn2-CH, consistent with Dyn2 severing emerging SNX1 tubules into vesicles. Indeed, FRAP of the lipid probe showed little or no recovery in the puncta, although the surrounding SLB recovered (Fig. 3D). The puncta thereforerepresent severed vesicles that are disconnected from the bilayer. Dyn2 thus converts SNX1-driven tubulation into vesicle formation in a GTP-dependent manner, which matches the longer SNX1 tubules in Dyn2 KO cells.

**Fig. 3.**
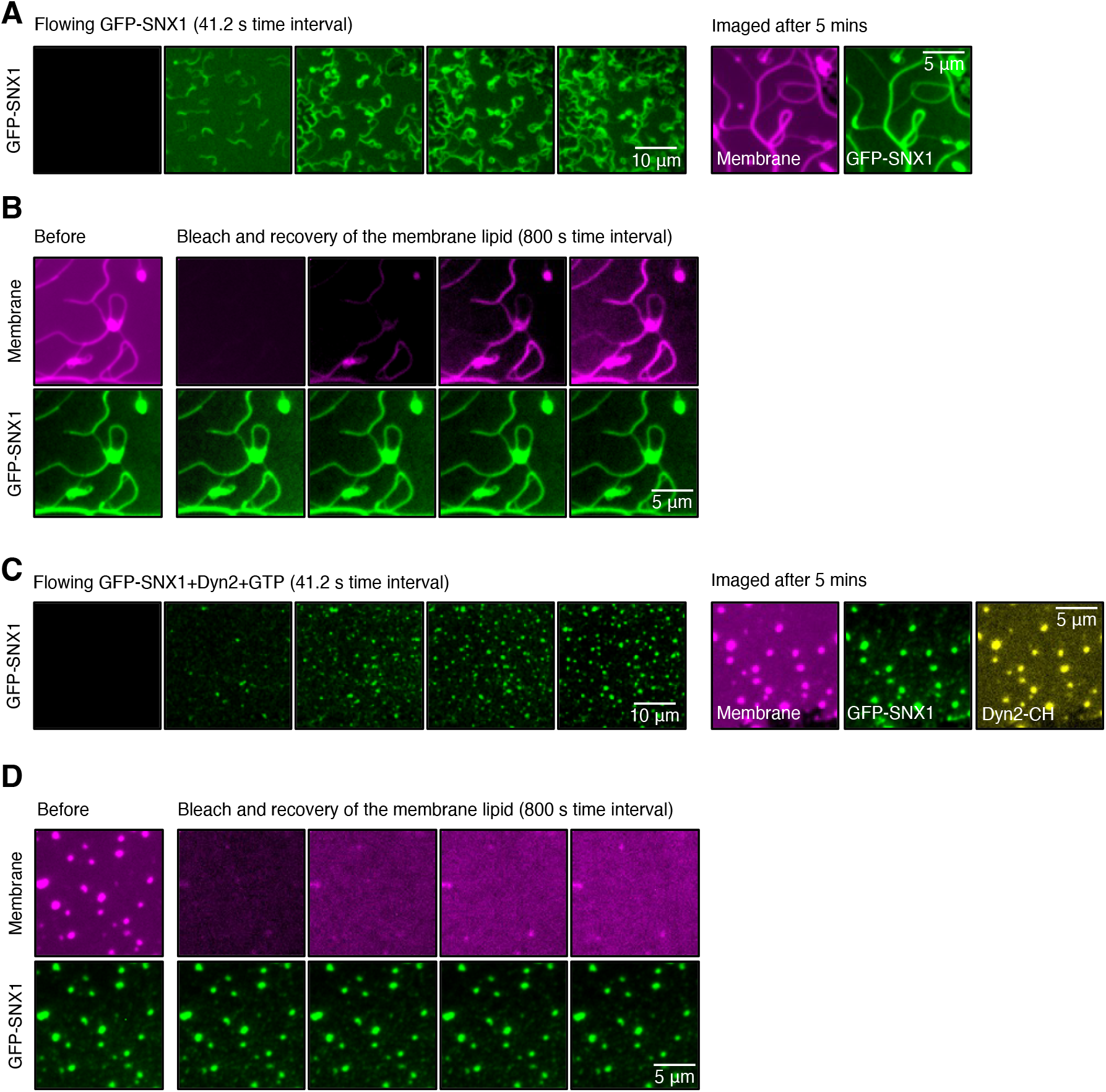
Dyn2 catalyzes fission of tubules formed by SNX1 alone. (A) Time-lapse montage (41.2 s interval) of GFP-SNX1 (green) flowed onto a supported lipid bilayer (SLB), showing progressive tubulation and formation of an extensive tubule network. The right panels show the membrane (magenta) and GFP-SNX1 (green) imaged after 5 min, confirming that the SNX1-positive structures are membrane tubules.Representative micrographs are shown from two independent experiments. (B) Fluorescence recovery after photobleaching (FRAP) of the lipid probe in the SLB after GFP-SNX1 addition. The membrane (top, magenta) and GFP-SNX1 (bottom, green) are shown before bleaching (left) and at successive time points after bleaching (right; 800 s interval). The lipid probe signal recovers, indicating that the tubules are continuous with the surrounding bilayer. Representative micrographs are shown from two experiments. (C) Time-lapse montage (41.2 s interval) of GFP-SNX1 flowed onto an SLB together with Dyn2 and GTP. Instead of tubules, GFP-SNX1 accumulates in discrete puncta that increase in number over time. The right panels show the membrane (magenta), GFP-SNX1 (green) and Dyn2-CH (yellow) imaged after 5 min. The puncta contain membrane, GFP-SNX1 and Dyn2, consistent with severing of emergent SNX1 tubules into vesicles. Representative micrographs are shown from two independent bilayers from one experiment. (D) FRAP of the lipid probe in the SLB after addition of GFP-SNX1, Dyn2 and GTP, imaged as in B. The puncta show little or no recovery, whereas the surrounding SLB recovers, indicating that the puncta are disconnected from the bilayer and represent severed vesicles. Representative micrographs are shown from one experiment.

### Loss of Dyn2 disrupts GLUT1 trafficking and reduces glucose uptake

Retromer–SNX1 tubules recycle cargo such as GLUT1 from endosomes to the plasma membrane, and we expect that failure to sever these tubules impairs recycling. We therefore examined GLUT1-GFP together with the lysosomal marker LAMP1-CH. In WT cells, GLUT1-GFP localized largely to the plasma membrane and overlapped only weakly with LAMP1-CH. In Dyn2 KO cells, GLUT1-GFP was largely absent from the plasma membrane and instead accumulated in a perinuclear compartment that overlapped with LAMP1-CH (Fig. 4A; orthogonal views). Colocalization analysis confirmed this shift (Fig. 4B). The Pearson’s correlation between GLUT1-GFP and LAMP1-CH was higher in Dyn2 KO cells than in WT cells (mean of the experiment medians ± SD, 0.775 ± 0.101 versus 0.480 ± 0.005; P < 0.0001, two-sided Mann-Whitney U test; n ≥ 50 cells per condition from at least three independent experiments). We then asked whether this mislocalization affects glucose uptake. Flow cytometry showed that Dyn2 KO cells took up 2.6-fold less of the fluorescent glucose analog 2-NBDG over 30 min than WT cells (Fig. 4C) (P < 0.0001, two-sided Mann-Whitney U test; n ≥ 25,000 cells per condition from three independent experiments). These results indicate that loss of Dyn2 diverts GLUT1 from the plasma membrane to a lysosomal compartment and lowers cellular glucose uptake.

**Fig. 4.**
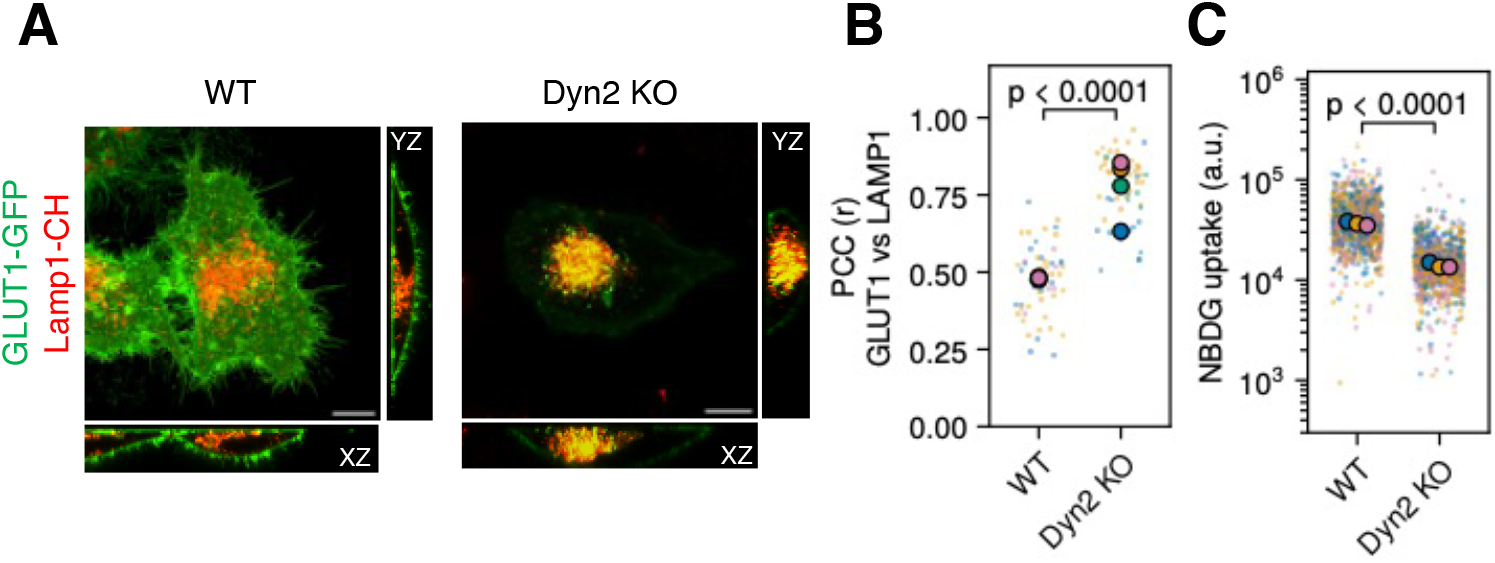
Loss of Dyn2 disrupts GLUT1 trafficking and lowers cellular glucose uptake. (A) Representative confocal images of GLUT1-GFP (green) in HeLa WT (left) and Dyn2 KO (right) cells; lysosomes are marked by LAMP1-CH (red). Main panels show MIPs, with orthogonal XZ and YZ views. GLUT1-GFP localizes largely to the plasma membrane in WT cells but overlaps with LAMP1-CH in a perinuclear compartment in Dyn2 KO cells. Scale bar, 10 µm. (B) Colocalization of GLUT1-GFP and LAMP1-CH, quantified by Pearson’s correlation coefficient (r). Small dots represent individual cells, and large dots represent the median of each independent experiment (color-coded by experiment). n ≥ 50 cells per condition from at least three independent experiments. A two-sided Mann-Whitney U test was used for statistical comparison. (C) Flow cytometry-based quantitation of 2-NBDG uptake (arbitrary units, log scale) by WT and Dyn2 KO cells after 30 min incubation with 100 µM 2-NBDG. Small dots represent individual cells, and large dots represent the median of each independent experiment (color-coded by experiment). n ≥ 25,000 cells per condition from three independent experiments. A two-sided Mann-Whitney U test was used for statistical comparison.

## Discussion

Our data identify Dyn2 as a fission factor for SNX1-coated endosomal tubules. A subset of Dyn2 puncta lie along SNX1 tubules, and its loss lengthens them and produces interconnected networks. Dyn2 binds SNX1 directly, an interaction that depends largely on its proline-rich domain (PRD) and requires the SNX1 N-terminus. With only SNX1, Dyn2 and GTP, tubules are converted into vesicles. Loss of Dyn2 diverts GLUT1 to a LAMP1-positive compartment and lowers glucose uptake.

SNX-BAR proteins are well established as the machinery that shapes endosomal tubules (Burd and Cullen, 2014; Cullen and Steinberg, 2018), but the factor that releases the resulting carriers has remained unclear (Cullen and Steinberg, 2018). A role for dynamin on mammalian endosomes was proposed long ago. Dominant-negative dynamin-K44A caused CI-MPR-containing tubules to accumulate, leading to the conclusion that dynamin mediates final scission (Nicoziani et al., 2000). Depletion of WASH or treatment with dynasore both produced long tubules pulled along microtubules, and WASH co-precipitated with dynamin-2, placing dynamin downstream of WASH (Derivery et al., 2009). In this model, actin pushes and microtubule motors pull, and the resulting tension facilitates dynamin-mediated fission. The WASH–Arp2/3 machinery was also linked to retromer-dependent sorting and scission (Gomez and Billadeau, 2009). However, these studies relied on dominant-negative mutants or dynasore, and none connected dynamin to a specific tubule-forming coat. Our findings address this gap - Dyn2 knockout lengthens SNX1 tubules, Dyn2 binds SNX1 directly, and Dyn2 is sufficient to sever SNX1-generated tubules in vitro. This reconstitution follows a two-component module we characterized previously. On SLBs, BIN1 - the BAR-domain containing protein that binds Dyn2, alone formed tubules, whereas BIN1 with Dyn2 and GTP produced tubules that were severed, leaving foci containing both proteins (Bhattacharyya and Pucadyil, 2024).

SNX1 behaves similarly, suggesting that a tubulating coat together with Dyn2 is sufficient for fission, consistent with the conceptual framework classifying Dyn2 as a specialized vesicle-fission dynamin tailored for narrow tubular substrates (Sarkar and Pucadyil, 2025). The yeast literature points to the same architecture. Vps1 localizes to endosomes decorated with retromer SNX-BARs and Mvp1 (the yeast SNX8 homolog), and its loss expands the SNX-BAR-coated network, lengthens tubule lifetimes two- to threefold, and sends Vps10 to the vacuole (Chi et al., 2014). Mvp1 binds Vps1 directly and recruits it to sever recycling tubules (Suzuki et al., 2021). Vps1 alone constricts and severs phosphoinositide-containing tubules in a GTP-dependent manner, and mutants unable to assemble or hydrolyze GTP fail to sever in vitro and mis-sort cargo in vivo (Gopan et al., 2026). Because yeast lacks WASH and spastin, and actin mutants show no sorting defect, actin is unlikely to be the major fission mechanism there (Chi et al., 2014). Dynamin-mediated scission may therefore be the ancestral core activity, with actin and other factors layered on in metazoans.

How Dyn2 engages SNX1 is of particular interest because the disordered SNX1 N-terminus is already a hub for assembly. Acidic Asp-Leu-Phe (DLF) motifs in the SNX1/2 N-terminus bind the SNX27 FERM domain (Yong et al., 2021; Simonetti et al., 2022), the same region binds VPS29 weakly, and the disordered N-termini of yeast SNX-BARs also determine complex specificity (Shortill, Frier and Conibear, 2022). We add Dyn2 as a binder of this region, although whether it competes with SNX27 and VPS29 or binds alongside them is yet to be determined. In yeast, an N-terminally stabilized Mvp1 tetramer blocks membrane binding, and this N-terminus is also needed for sorting (Shortill, Frier and Conibear, 2022). Sequential or competitive use of a flexible N-terminus could therefore ensure that scission machinery is recruited only to mature tubules. The PRD side of the interaction is also unusual. Dynamin PRDs normally bind SH3 domains, and efficient endocytosis depends on several SH3 domains engaging adjacent PRD motifs (Rosendale et al., 2019; Bhattacharyya and Pucadyil, 2024), yet SNX1 has no SH3 domain.

Dyn2 is unlikely to be the only factor acting on these tubules. Dynein–dynactin provides pulling force, and a constriction factor such as dynamin was suggested to assist release (Wassmer et al., 2009). LITAF and CDIP1 partition into SNX1-labeled tubules, and their loss expands tubular recycling compartments (Wunderley et al., 2021). Because Dyn2 is sufficient in vitro, we need not treat these as obligatory partners. Instead, they may set the timing, position or efficiency of scission in cells. Cytoskeletal forces would raise membrane tension, which favors dynamin (Derivery et al., 2009), while LITAF might act on a parallel subset of tubules.

The GLUT1 phenotype places Dyn2 within known sorting pathways. GLUT1 is a well-characterized SNX27–retromer cargo. In SNX27 or VPS35 knockout cells GLUT1 is mis-sorted to LAMP1-positive lysosomes and degraded faster (Steinberg et al., 2013; Simonetti et al., 2022), and mutating its PDZ-binding motif has the same effect and reduces glucose uptake (Wieman et al., 2009). Surface delivery is further regulated by autophagy through TBC1D5 (Roy et al., 2017) and by PTEN (Shinde and Maddika, 2017). Dyn2 KO cells show the same pattern of lysosomal diversion and reduced glucose uptake, as does loss of Vps1 in yeast (Chi et al., 2014; Gopan et al., 2026). This outcome is not the only one possible, and the distinction matters. Co-depletion of SNX1/2 or SNX5/6 shifted GLUT1 to EEA1-positive endosomes without increased lysosomal degradation (Steinberg et al., 2013), and disrupting the SNX27 FERM–SNX1 interface blocks surface delivery without impairing retrieval from lysosomes (Yong et al., 2021; Simonetti et al., 2022). In contrast, our Dyn2 KO cells accumulate GLUT1 in a LAMP1-positive compartment, resembling SNX27/VPS35 loss more than SNX-BAR loss. Chronic loss of scission might allow cargo stuck in unresolved tubules to mature into the degradative pathway. Alternatively, Dyn2 may act at more than one step, since it associates with WASH, part of the SNX27–retromer assembly (Derivery et al., 2009; Guo et al., 2024), and has other cellular roles.

If Dyn2 is a general scission factor for SNX1 tubules, its effects should extend beyond GLUT1. SNX1/2–SNX5/6 heterodimers (ESCPE-1) mediate retrograde CI-MPR transport independently of the retromer trimer (Kvainickas et al., 2017; Simonetti et al., 2017), and CI-MPR is where dynamin involvement was first suggested (Nicoziani et al., 2000). CI-MPR trafficking and cathepsin delivery should therefore be impaired in Dyn2 KO cells. SNX-BARs also act in SNX3-dependent Wntless transport (Harterink et al., 2011) and in EGFR sorting with SNX1 and SNX32 (Wang et al., 2024), both of which would be worth testing. SNX1 tubules are also conserved in plants, where Arabidopsis SNX1 knockdown mis-sorts VSR2 (Li et al., 2026), suggesting that tubule scission is a feature of SNX1 pathways across kingdoms.

Dyn2 also functions in clathrin-mediated endocytosis, but a primary internalization defect would be expected to retain GLUT1 at the plasma membrane. Instead, we observe loss of peripheral GLUT1 and its accumulation in a LAMP1-positive compartment, consistent with a post-endocytic recycling defect. Because Dyn2 loss is chronic, indirect effects on endosomal organization through other Dyn2-dependent processes remain possible. Finally, the SLB assay shows that Dyn2 is sufficient to sever SNX1 tubules without WASH or other dynamin isoforms, but whether this is the dominant route in cells remains to be determined. These findings may also be relevant to disease. VPS35 mutations cause Parkinson disease (Vilariño-Güell et al., 2011; Zimprich et al., 2011), and retromer dysfunction is linked to Alzheimer disease and other neurological disorders (Small et al., 2005; Small and Petsko, 2015). A DNAJC13 variant alters SNX1-enriched tubules in pharmacogenetic Parkinson models (Yoshida et al., 2024), so how SNX1 carriers are cut may bear on these mechanisms. Dyn2 mutations are likewise associated with centronuclear myopathy (Bhattacharyya and Pucadyil, 2024), and they may affect whether endosomal tubules deliver cargo with fidelity. Finally, retromer and sorting nexins are exploited by pathogens including Legionella and HPV16 (Finsel et al., 2013; Popa et al., 2015; Lu et al., 2022), making a defined fission step a plausible point of vulnerability.

### Materials and Methods Cell culture

HeLa cells were maintained in Dulbecco’s modified Eagle’s medium (DMEM) supplemented with 10% (v/v) fetal bovine serum (FBS) and 1% (v/v) penicillin-streptomycin (HiMedia) at 37°C in a humidified atmosphere with 5% CO_2_

### Generation of Dyn2 knockout and stable GFP-SNX1 cell lines

To generate Dyn2 knockout cells, gRNAs targeting two distinct exons of Dyn2 (gRNA 1: 5′-GTGACGATTCCTGAACCGCG-3′; gRNA 2: 5′-TTGACGAGAAGGACTTACGA-3′) were each cloned into the pSpCas9(BB)-2A-GFP (PX458) backbone (Addgene plasmid #48138), and the two plasmids were co-transfected into cells. GFP-positive cells were sorted by FACS into single cells to establish clonal lines, and loss of Dyn2 in the resulting clones was confirmed by western blot. WT and Dyn2 KO cells stably expressing GFP-SNX1, or GFP alone as a control, were generated by lentiviral transduction with GFP-SNX1 or GFP cloned into the pLVX backbone, followed by FACS enrichment. WT and Dyn2 KO cell populations were sorted to obtain comparable, low levels of GFP-SNX1 expression.

### Imaging GFP-SNX1 and Dyn2

Cells stably expressing low levels of GFP-SNX1 were seeded on glass-bottom dishes coated with poly-D-lysine (Gibco) 48 h before the experiment. Cells were fixed with 4% (w/v) paraformaldehyde in DPBS (pH 7.4), permeabilized with 0.1% (w/v) Triton X-100, and blocked with 1% BSA in DPBS containing 0.1% Tween-20. Cells were then incubated overnight at 4°C with an anti-Dyn2 antibody (Abcam, ab65556, 1 µg/ml), washed, and incubated with an Alexa Fluor 647-conjugated anti-rabbit IgG secondary antibody (Invitrogen). Cells were imaged on an Olympus FLUOVIEW FV4000 laser-scanning confocal microscope (software v3.2.1.85) mounted on an inverted IX83 stand, using a UPLXAPO 60× oil-immersion objective (NA 1.42, working distance 0.15 mm, immersion refractive index 1.518). EGFP was excited with the 488 nm diode laser and Alexa Fluor 647 with the 640 nm diode laser, using a DM405/488/561/640/730 excitation dichroic. Fluorescence was collected sequentially (frame-wise) on two spectral detectors, at 495–535 nm for EGFP and 650–710 nm for Alexa Fluor 647. The pinhole was set automatically to 231 µm. Images were collected with a galvanometer scanner at zoom 1.4, as 1024 × 1024 pixel, 16-bit frames (0.148 µm/pixel), with a 2 µs pixel dwell time (unidirectional scanning) and 4× line averaging.Z-stacks comprised 18 optical sections at 0.5 µm spacing (8.5 µm total range). Laser transmissivity was 5% (488 nm) and 10% (640 nm).

### GFP-SNX1 imaging for tubule length quantitation

Live cells stably expressing GFP-SNX1 were imaged in HEPES-buffered Hanks’ balanced salt solution (HBSS). Images were acquired on an Olympus FLUOVIEW FV4000 laser-scanning confocal microscope under settings described above, but with the 488 nm laser transmissivity at 2%. Maximum intensity projections (MIPs) were generated from raw z-stacks, and individual cells were cropped for downstream analysis. To segment the GFP-SNX1–positive compartment from background, a pixel classification model was trained in ilastik (Berg et al., 2019) using an annotated training set of 50 representative single-cell images. The resulting probability maps were thresholded to generate binary masks across all single-cell datasets. Tubule morphology was quantified from these binary images using the Mitochondria Analyzer plugin for Fiji (Chaudhry, Shi and Luciani, 2020) to extract individual branch lengths per cell. For comparative analyses, the maximum tubule length per cell was identified and plotted.

### Co-immunoprecipitation and western blotting

Cells stably expressing GFP-SNX1 or GFP alone (control) were grown to ∼90% confluency in two 10-cm dishes. Cells were harvested and lysed in 800 µl of ice-cold lysis buffer (50 mM Tris-HCl, pH 7.5, 150 mM NaCl, 5 mM MgCl_2_, 5% glycerol, 0.5% NP-40, 1 mM PMSF and 1× protease inhibitor cocktail). Lysates were clarified by centrifugation at 12,000 × g for 10 min at 4°C, and an aliquot (1%) was set aside as the input control. The remaining lysate was incubated for 2 h at 4°C with rotation with Dynabeads (Thermo Fisher Scientific) pre-bound to anti-GFP antibody (Invitrogen, MA5-15256; 1:200). Beads were washed three times with wash buffer (50 mM Tris-HCl, pH 7.5, 150 mM NaCl, 5 mM MgCl2, 0.1% NP-40), and bound complexes were eluted by boiling in 2× Laemmli sample buffer for 10 min. Eluates and input samples were resolved on 10% SDS-PAGE gels and transferred to PVDF membranes. Membranes were blocked with 5% BSA in 1× TBS-T for 1 h at room temperature (RT) and incubated for 3 h at RT with the following primary antibodies: anti-GFP (Invitrogen, MA5-15256; 1:5000) to confirm immunoprecipitation, anti-Dyn2 (Abcam, ab65556; 1:3000) to assess co-precipitation, and anti-β-actin (Santa Cruz, sc-47778; 1:5000) as a loading control for inputs and whole-cell lysates. After three 5-min washes with 1× TBS-T, membranes were incubated with the appropriate HRP-conjugated secondary antibodies for 1 h at RT, followed by three further 5-min washes with 1× TBS-T. Signals were developed using an ECL chemiluminescent substrate (Thermo Fisher Scientific) and imaged on an iBright 1500 imaging system (Thermo Fisher Scientific). Immunoblots were processed using Fiji (Schindelin et al., 2012).

### Expression and purification of proteins

For bacterial expression, the following constructs were cloned into pET15b: 6xHis-TEV-GFP-SNX1-StrepII, 6xHis-TEV-Dyn2-mCherry(CH)-StrepII, 6xHis-TEV-GFP, 6xHis-TEV-CH, 6xHis-TEV-GFP-SNX1ΔN (Δ1-142)-StrepII, and 6xHis-TEV-Dyn2ΔPRD (Δ744-870)-CH-StrepII. Human SNX1 and Dyn2 were used. Recombinant proteins were expressed in T7 Express cells (NEB). Protein expression was induced with 0.5 mM IPTG at an OD600 of 0.6-0.8 and grown at 18°C for 12-14 h. Cells were harvested, and pellets were stored at −40°C until use. Pellets were resuspended in lysis buffer (20 mM HEPES, pH 7.6, 500 mM NaCl), lysed by sonication, and clarified by centrifugation at 30,000 × g. The soluble fraction was incubated with TALON resin (Takara) for 1 h at 4°C. For 6xHis-StrepII dual-tagged proteins, the resin was transferred to a PD-10 column, washed with 100 mL of lysis buffer, and eluted with 50 mM EDTA in lysis buffer. The eluate was loaded onto a 5 mL StrepTactin HP column (Cytiva), washed with assay buffer (20 mM HEPES, pH 7.6, 150 mM NaCl), and eluted with 2.5 mM desthiobiotin (Sigma) in assay buffer. For 6xHis-only proteins, the TALON resin was washed sequentially with lysis buffer and assay buffer, and proteins were eluted with 50 mM EDTA in assay buffer. Where required, eluates were dialyzed overnight against assay buffer at 4°C. Before use in assays, all proteins were centrifuged at 100,000 × g for 30 mins at 4°C to remove aggregates.

### Bead interaction assay, imaging and image analysis

Purified 6xHis-tagged bait proteins (1 µM) were immobilized on TALON metal affinity resin (Takara; 50 µL settled resin in a total volume of 1mL) for 30 min at 4°C and washed three times with assay buffer. 6xHis-tagged prey proteins were incubated overnight at 4°C with 6xHis-TEV protease (10:1 protein-to-protease molar ratio) in assay buffer supplemented with 1 mM DTT. The reaction mixture was dialyzed against assay buffer to remove DTT and passed back over TALON resin to deplete uncleaved prey and 6xHis-TEV; the flow-through containing cleaved prey was collected. Cleaved prey (2 µM in 1mL) was incubated with bait-bound beads for 30 min in assay buffer containing 20 mM imidazole to prevent non-specific binding. Beads were washed with the assay buffer with 20 mM imidazole to remove unbound protein and added to a lab-tek chamber. Beads were imaged on an Olympus IX83 inverted microscope with a 20× objective (UPLSAPO20X) and a Photometrics Evolve 512 EMCCD camera (frame-transfer mode, 10 MHz, 16-bit readout, 1×1 binning, 100 ms exposure) using Micro-Manager 1.4.21. Images were acquired using a pE4000 (CoolLED) light source sequentially as 512 × 512, 16-bit images, with settings held constant within each experiment. Beads were segmented independently in each field from a single mask channel (FITC for GFP-bait series, Texas Red for CH-bait series) by Gaussian smoothing (σ = 1 px) and Otsu thresholding, followed by hole filling and removal of objects ≤200 px as debris. All channels were quantified within each bead’s mask, regardless of the segmentation channel. Local background was the median intensity in an annulus 10–25 px outside the bead edge, excluding a 10-px dilation of all segmented beads in the field, and thus also absorbs camera offset and diffuse signal from unbound protein. The whole-bead mean intensity minus its local background was estimated. GFP signals were normalized to the mean FITC signal of GFP-bait, no-prey beads, and CH signals to the mean Texas Red signal of CH-bait, no-prey beads. Images were rendered using Fiji (Schindelin et al., 2012).

### Supported Membrane Templates (SMrT) generation and imaging

SMrTs were generated as described previously (Dar, Kamerkar and Pucadyil, 2015; Dar, Kamerkar and Pucadyil, 2017; Bhattacharyya and Pucadyil, 2024; Swaminathan and Pucadyil, 2024). A lipid mixture of 1,2-dioleoyl-sn-glycero-3-phospho-L-serine (DOPS), 1,2-dioleoyl-sn-glycero-3-phosphocholine (DOPC), 1,2-dioleoyl-sn-glycero-3-phospho-(1’-myo-inositol-3’-phosphate) (PI(3)P; Avanti Polar Lipids), and 1,1’-dioctadecyl-3,3,3’,3’-tetramethylindodicarbocyanine 4-chlorobenzenesulfonate (DiD; Invitrogen) was prepared in chloroform at a 15:79:5:1 molar ratio and a total lipid concentration of 1 mM. Using a glass syringe, 2 µL of this solution was deposited onto a PEGylated (PEG400) glass coverslip. After the solvent evaporated, the coverslip was mounted in an FCS2 flow chamber (Bioptechs). SMrTs were formed by filling the chamber with assay buffer at a high flow rate. 100 nM GFP-SNX1 in the absence or presence of 200 nM Dyn2-CH with 1 mM GTP (Jena Bioscience) and 1 mM MgCl_2_ were flowed onto SMrTs in assay buffer at low flow rates and imaged on an Olympus IX83 inverted microscope with a UPLSAPO 100× oil-immersion objective and a Photometrics Evolve 512 EMCCD camera (frame-transfer mode, 10 MHz, 16-bit readout, multiplication gain 10, 1×1 binning, 100 ms exposure) using Micro-Manager 1.4.21. Images were acquired using a pE4000 (CoolLED) light source sequentially in the FITC (GFP) and Texas Red (CH) and Cy5 (DiD) channels as 512 × 512 pixel, 16-bit images, with settings held constant within each experiment. For wide-field fluorescence recovery after photobleaching (FRAP), supported lipid bilayers were incubated with the indicated proteins and washed to remove unbound protein. A wide area of the membrane containing the lipid probe (DiD) was photobleached with a brief pulse of high-intensity light. Illumination was then reduced, and recovery was recorded at 10-s intervals. Images were rendered using Fiji (Schindelin et al., 2012).

### GLUT1 and LAMP1-CH imaging

WT and Dyn2 KO cells were co-transfected with GLUT1-GFP (Addgene plasmid #18729) and LAMP1-CH (Addgene plasmid #55073). After transfection, cells were maintained in DMEM containing 1 g/L glucose, supplemented with 10% FBS and 1% (v/v) penicillin-streptomycin. At 48 h post-transfection, cells were treated with 100 nM bafilomycin A1 (Cell Signaling Technology) for 4 h at 37°C to reduce lysosomal acidification and thereby allow detection of GFP fluorescence. Cells were then imaged live at 37°C in HBSS on an Olympus FLUOVIEW FV4000 laser-scanning confocal microscope (software v3.2.1.85) mounted on an inverted IX83 stand, using a UPLXAPO 60× oil-immersion objective (NA 1.42, working distance 0.15 mm). EGFP was excited with the 488 nm laser and mCherry with the 561 nm laser, using a DM405/488/561/640/730 excitation dichroic. Emission was collected on two spectral detectors, at 500–540 nm (EGFP) and 570–670 nm (mCherry), with the pinhole set automatically to 219 µm. Channels were acquired in line-sequential mode with a galvanometer scanner at zoom 1.4, as 1024 × 1024 pixel, 16-bit images (0.148 µm/pixel), with a 2 µs pixel dwell time (unidirectional scanning) and 2× line averaging. Z-stacks comprised 21 optical sections at 0.5 µm spacing (10 µm total range). Laser transmissivity was 5% for both the 488 nm and 561 nm lines. Colocalization between GLUT1-GFP and LAMP1-CH was quantified using Huygens.

### 2-NBDG uptake assay

Glucose uptake in WT and Dyn2 KO cells was measured using the fluorescent glucose analogue 2-NBDG (Invitrogen). Cells were starved in glucose-free medium for 20 min at 37°C and then incubated with 100 µM 2-NBDG for 30 min at 37°C. Cells were rinsed with ice-cold DPBS, detached with trypsin, pelleted, and resuspended in DPBS. Samples were analyzed immediately on a BD flow cytometer controlled by BD FACSDiva software (v8.0.1.1). GFP/FITC fluorescence was excited by the 488 nm laser and collected in the FITC channel. Cells were first gated on FSC-A versus SSC-A (P1) to exclude debris, and single cells were then selected from P1 by FSC-A versus FSC-H (P2). 2-NBDG fluorescence intensity was measured in the P2 population. Data were analyzed using Floreada.io.

### Statistical analysis and data rendering

Statistical analysis and data rendering were performed using custom scripts written using Python 3.15.0.

## Acknowledgements

M.S. thanks the Council of Scientific & Industrial Research for a graduate fellowship. S.D. thanks the Department of Biotechnology (DBT) for a graduate fellowship. T.J.P. thanks the DBT/Wellcome Trust India Alliance for a Team Science Grant (Grant No. IA/TSG/21/1/600245), the Anusandhan National Research Foundation for a SUPRA Grant (Grant No. SPR12021100014) and the Howard Hughes Medical Institute for an International Research Scholar’s Grant (Grant No. 55008746). We thank Sunando Dutta (IISER Bhopal, India) for the SNX1 construct.

